# Survival strategies of *Enterococcus mundtii* in the gut of *Spodoptera littoralis*: a live report

**DOI:** 10.1101/2020.02.03.932053

**Authors:** Tilottama Mazumdar, Beng Soon Teh, Aishwarya Murali, Wolfgang Schmidt-Heck, Yvonne Schlenker, Heiko Vogel, Wilhelm Boland

**Author notes:** Address correspondence to: Tilottama Mazumdar, Wilhelm Boland.

## Abstract

The complex interaction between a higher organism and its resident gut flora is a subject of immense interest in the field of symbiosis. Many insects harbor a complex community of microorganisms in their gut. Larvae of *Spodoptera littoralis*, a lepidopteran pest which is prevalent in tropical and subtropical regions of the world, have a tube-like gut structure containing a simple bacterial community. This community varies both spatially (along the length of the gut) and temporally (during the life cycle of the insect).

To monitor the dynamics and rapid adaptation of microbes to the gut conditions, a GFP-tagged reporter *E. mundtii* was constructed. After feeding to early instar *S. littoralis* larvae, the tagged-microbes recovered from the fore and hind guts by flow cytometry. The fluorescent reporter confirmed the persistence of *E. mundtii* in the gut. RNA-sequencing of the sorted bacteria highlighted various strategies that the symbiont employs to survive, including upregulated pathways for tolerating alkaline stress, forming biofilms and two-component signaling systems, resisting oxidative stress and quorum sensing. Although these symbionts depend on the host for amino acid and fatty acids, differential regulation among various metabolic pathways points to an enriched lysine synthesis pathway in the hindgut of the larvae.

## Introduction

Insects comprise the largest phylum of arthropods, according to the IUCN red list. Microorganisms have been known to form symbiotic relationships with insects by supplying the latter with essential nutrients, protection against pathogens, and aiding in digesting recalcitrant organic matter. They contribute significantly to insects’ ability to act as potential pathogens to animals, pests or pollinators of food crops, and as cyclers of carbon and nitrogen during the decomposition of plant biomass (1)

There are several factors that determine the gut bacterial composition in insects. The gut can be compartmentalized, resulting in structures that vary according to the complexity of the microbial communities. Insects with a straight, tube-like gut usually possess a less diverse microbial population compared to species with invaginations and deep pouches (1). Other factors that shape the gut population include the following: oxygen level, gut pH, the presence of digestive enzymes, antimicrobial compounds, and insect diet (2, 3). Although most bacteria have an affinity for neutral pH, several acidophiles and alkalophiles have adapted to extreme pH conditions.

Gut microbes can be either vertically or horizontally transmitted. Vertical transmission allows bacterial transfer (from the ovaries to the egg shells) to the next generation (4), whereas horizontal transmission occurs over the course of the life cycle, through diet and social behavior.

Regardless of how bacteria are transmitted, microbial populations may be unstable during early developmental stages (5, 6). For example, in holometabolous insects, a complete metamorphosis of the gut occurs in the larva, through pupal and adult stages, resulting in microbial turnover and variable microbial counts (5).

Insects are helped by their bacterial and fungal symbionts with functions relating to the digestion of complex plant carbohydrates and amino acids, the assimilation of vitamins and the development of defensive strategies against plants and animals (7), (8), (9). Higher animals employ their resident gut symbionts for digestion of recalcitrant substrates. The plant cell-wall component pectin is one example of a polymer, which in the leaf beetle *Cassida rubiginosa* digests by employing an extracellular bacterium (10). Likewise, termites depend on their gut bacteria, archaea and eukaryotic gut symbionts for the breakdown and fermentation of plant fiber to acetate and methane. The gut microbiota further help in digesting xylan, arabinogalactan and carboxymethylcellulose, and can even work as a bio-reactor capable of mineralizing humus and nitrogen cycling (11, 12). Since the phloem sap is low in vitamins, amino acids and nitrogen, aphids have to rely on their primary symbiont, *Buchnera aphidicola* for nutrition. House crickets are hosts for *Acheta domesticus* bacteria in their hind guts, enabling the insects to utilize plant polysaccharides (2).

In the course of evolution of plant-insect associations, insects have in several cases countered plant defensive strategies and managed to digest usually inaccessible, but highly nutritious plant polymers by acquiring several enzymes relating to digestion of complex plant carbohydrate through horizontal gene transfer (7). In other cases, functions relating to amino acid and vitamin assimilation (9), (13), and counter-defensive strategies against plants have been obtained by the insects from bacteria or fungi. For example, *Burkholderia gladioli* produce a mixture of antifungal antibiotics that protect the egg stages of Lagriinea beetles, which would otherwise be infected by the fungus *P. lilacinum* (14).

The cotton leafworm, *Spodopera littoralis* is a notorious pest feeding on a broad range of plants. Larvae of this species have a longitudinal gut structure without compartments, increasing the chances of flushing out bacteria, possibly preventing long-term colonization. This simple gut structure could be one reason for the overall low gut-bacterial density observed in Lepidoptera (1), (15). Despite the seemingly simple structure, a pH gradient exists along the gut: the anterior part and the midgut of lepidopteran larvae is highly alkaline, with a pH range of 11-12 (16), but is neutral in the posterior part (17), which restricts the survival of many microbial species. Despite the alkaline pH, bacteria of the phylum Fermicutes, notably Enterococci and *Clostridium* sp. have emerged as core gut bacteria in the larval stages of *Spodoptera littoralis* (5). In particular, the dominant *E. mundtii* has been shown to colonize the gut of *S. littoralis* throughout different developmental stages (3), (5), (18).

Enterococci are successful colonizers of the gastrointestinal tract of humans, animals, and insects. They can exert probiotic, positive effects which have been shown in humans (19). *Enterococcus mundtii* is a gram positive, non-motile lactic acid bacterium abundantly found in soil, human navel, cow teats and on hands of milkers. They are well adapted to dairy and plant environment (20)

Antimicrobial activity has been shown for several Enterococci species, particularly *E. mundtii* isolated from a lepidopteran insect. *E. mundtii* produce an antimicrobial peptide called mundticin KS, that keeps potential pathobionts like *Enterococcus fecalis* and *Enterococcus casseliflavus* at bay. These pathobionts are early colonizers apparent in the first instar larvae, followed by a steep reduction in their numbers, owing to mundticin (5), (3). Despite potential regulatory effects on the gut microbiome, the specific contributions of Enterococci in insects remain largely unknown. Larvae of several Lepidopteran species produce high concentrations of 8-hydroxyquinoline-2-carboxylic acid, an iron chelator that derived from tryptophan and found in their gut and regurgitate (21). Since iron is one of the main elements in several metabolic pathways, such as quenching of reactive oxygen species, oxidative metabolism in TCA cycle, electron transport chain, nitrogen assimilation and many more (22), this chelator is assumed to control the microbiome in larval guts.

In this paper we focused on how a particular bacterium’s quick and efficient adaptation processes to the changing living conditions from *in vitro* culture conditions in the laboratory, to the *Spodoptera littoralis* gut environment.

## Results

The GFP-tagged reporter *Enterococcus mundtii* (18) was allowed to colonize the gut environment of *Spodoptera littoralis* and was later recovered and sorted using flow cytometry to compare its gene expression to that of *E. mundtii* grown *in vitro*.

### 1. Flow cytometry

The reporter bacteria of *Enterococcus mundtii* which were exposed to the gut conditions of *Spodoptera littoralis* larvae were sorted and isolated using flow cytometry. Next, the pooled bacteria were used to study changes in their transcriptome, as compared to the same bacteria grown in Todd Hewitt Broth (THB). We chose THB-cultured *E. mundtii* grown in a shaker incubator at 37 degree Celsius and 220 RPM as a control because these are ideal, stress-free conditions. THB is a complete medium, in which bacteria can grow on dextrose as the source of energy. Since the pH in *S. littoralis* gut is alkaline in the foregut and neutral in the hindgut, we focused on the *E. mundtii* growing at the two terminals of the gut.

From the gut homogenates containing the fluorescent reporter *E. mundtii,* 250,000 fluorescent cells were sorted by a flow cytometer. The collected cells constituted 2 to 4% of the total homogenate. In comparison, 250,000 fluorescent *E. mundtii* cells grown *in vitro* were also sorted for our differential gene expression analysis (Fig. S1).

### 2. RNAseq, mapping and differential gene expression analysis

The RNA extracted from the FACS-sorted *E. mundtii* cells was sequenced using the Illumina Ultra-Low Input RNA kit and the resulting 10 million short reads per treatment and replicates were processed and aligned against the fully sequenced genome of *Enterococcus mundtii* QU25 (23). Overall, the three replicates of bacterial cells isolated from foregut, hindgut and control had varying levels of alignment percentages ranging from 17.5 to 55.5, as shown in supplementary table S2. Since the concentration of the RNA obtained from the FACS-sorted gut reporter *E. mundtii* was low, we performed total RNA amplification. To maintain uniformity, the control samples were treated the same way.

The numbers of up and down regulated genes between *E. mundtii* cells exposed to different *S. littoralis* gut sections is shown Fig. 1. 284 and 275 genes are significantly differentially regulated (fold change = 2, p ≤ 0.05) in *Enterococcus mundtii* in the fore and hind guts, respectively. Density plot in Fig. S3 shows the distribution of differentially expressed genes in foregut, hindgut and control.

**Figure 1:**
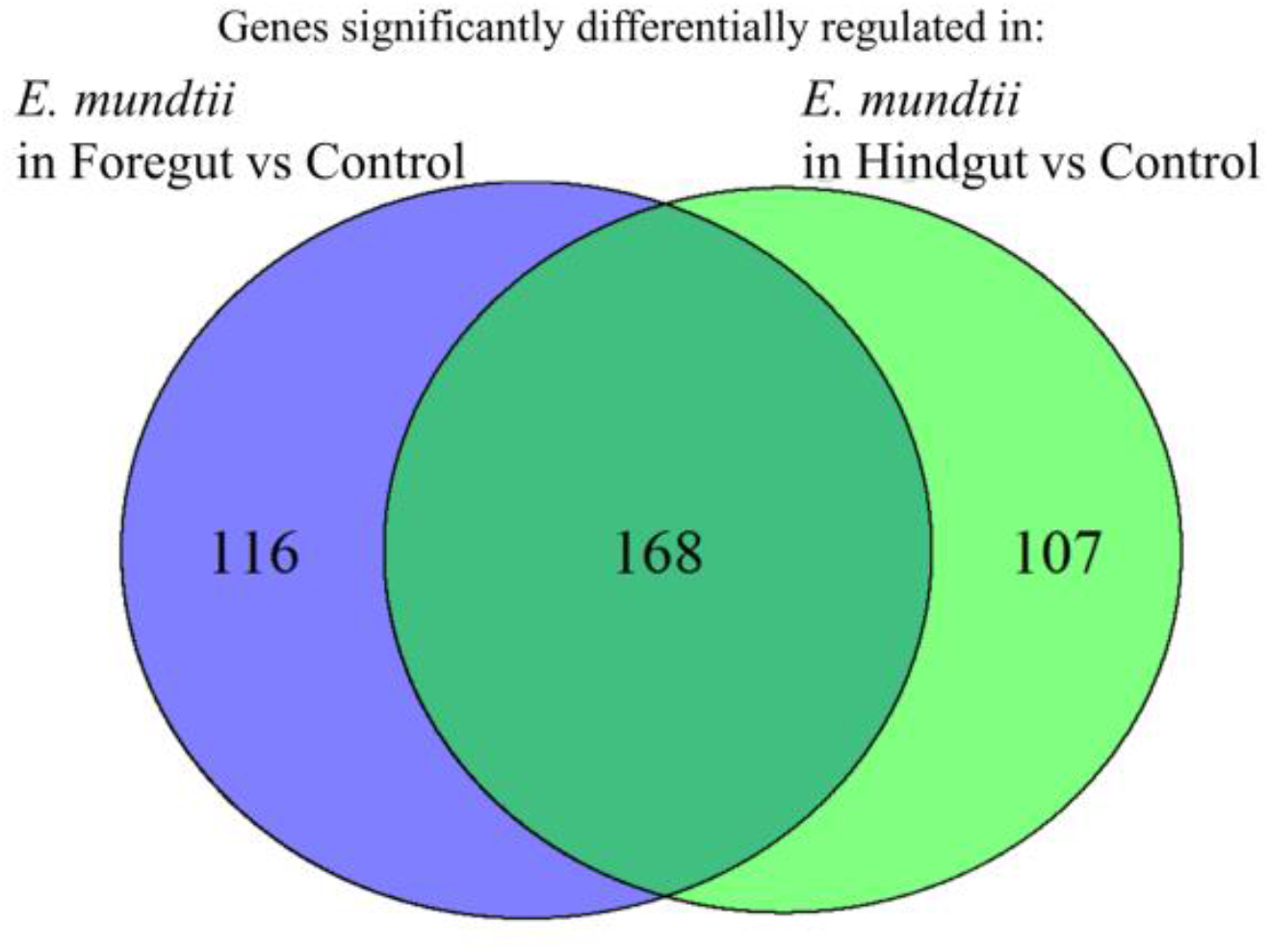
Venn diagram showing overlap of significantly differentially expressed genes (supplementary) in the following two conditions: *E. mundtii* living in foregut vs control, and *E. mundtii* living in hindgut vs control.

There are 169 genes in common between the *E. mundtii* exposed to the fore and hindguts that are differentially regulated when compared to the control. Most of these common genes belong to mechanisms required by *E. mundtii* to colonize by adhering to the gut wall, avoid stresses, and acquire iron and complex carbohydrates ((Fig. 1), (supplementary S7)).

To test for biological and technical variability, individual replicates were analyzed and a PCA plot (Fig. 2) and dendrogram (Fig. 3) were generated. The gene expression profiles of *E. mundtii* from the insect gut and the control form separate clusters and nodes, clearly separating them from each other.

**Table 1:**
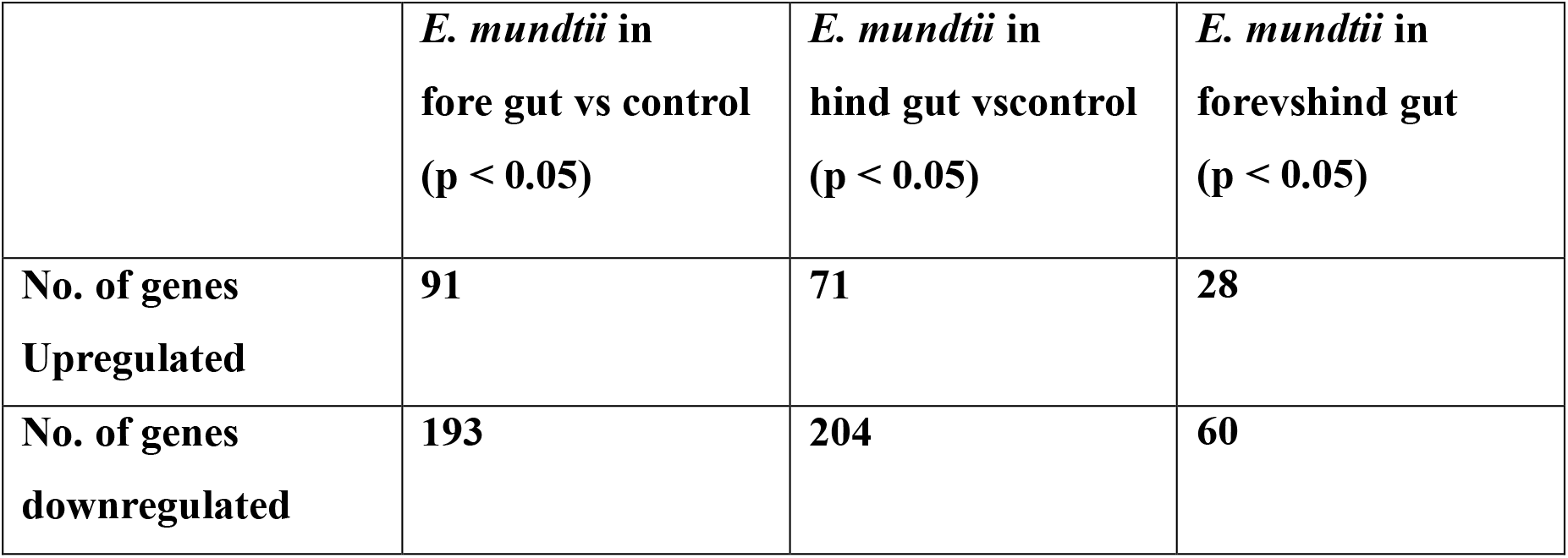
Table showing the numbers of significantly differentially expressed genes-up- and downregulated (p ≤ 0.05) in *Enterococcus mundtii*–by comparing the following conditions: *E. mundtii* living in foregut vs control, hindgut vs control and foregut vs hindgut.

|  | *E. mundtii* in<br>fore gut vs control<br>(p < 0.05) | *E. mundtii* in<br>hind gut vs control<br>(p < 0.05) | *E. mundtii* in<br>fore vs hind gut<br>(p < 0.05) |
| --- | --- | --- | --- |
| No. of genes<br>Upregulated | 91 | 71 | 28 |
| No. of genes<br>downregulated | 193 | 204 | 60 |

**Figure 2:**
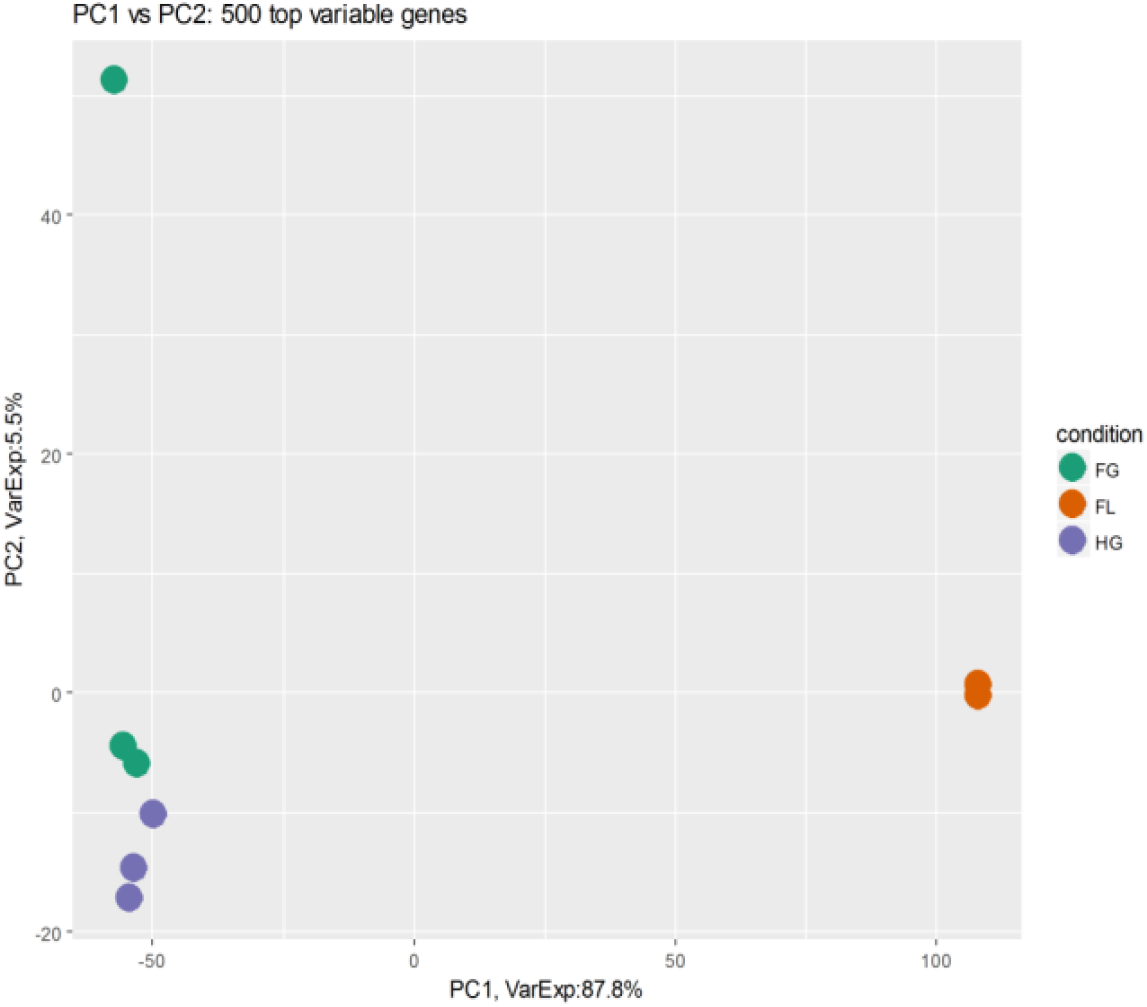
PCA plot showing clustering of the transcriptomic profiles among the three technical replicates of *E. mundtii* obtained from the foregut (FG), hindgut (HG) and control (FL).

**Figure 3:**
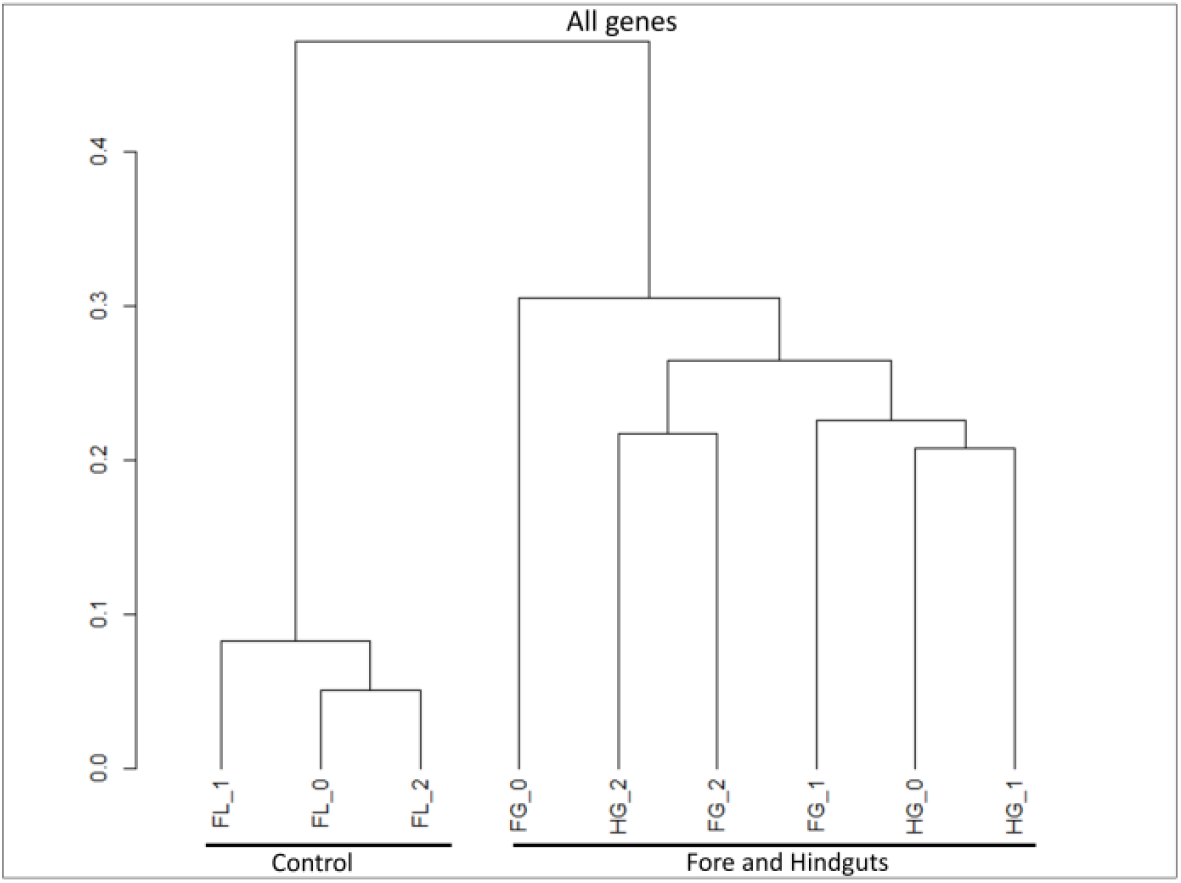
Dendrogram showing hierarchical clustering among the three replicates of transcriptomes analyzed from *E. mundtii* obtained from the foregut, hindgut and control. The gene expression profiles from the foreguts (FGs) and the hindguts (HGs) partially overlap and cluster away from the control (FL) profile.

### 3. Annotations based on gene ontology of proteins

The full picture of the pathways from *E. mundtii* living in the gut of *S. littoralis* is evident in Fig. 4. The differentially expressed genes have been classified according to the three categories of gene ontology: molecular function, biological process and cellular components. We only discuss the category of biological processes, which highlights the major pathways the bacteria are involved in when they are living in the gut of the host.

**Figure 4:**
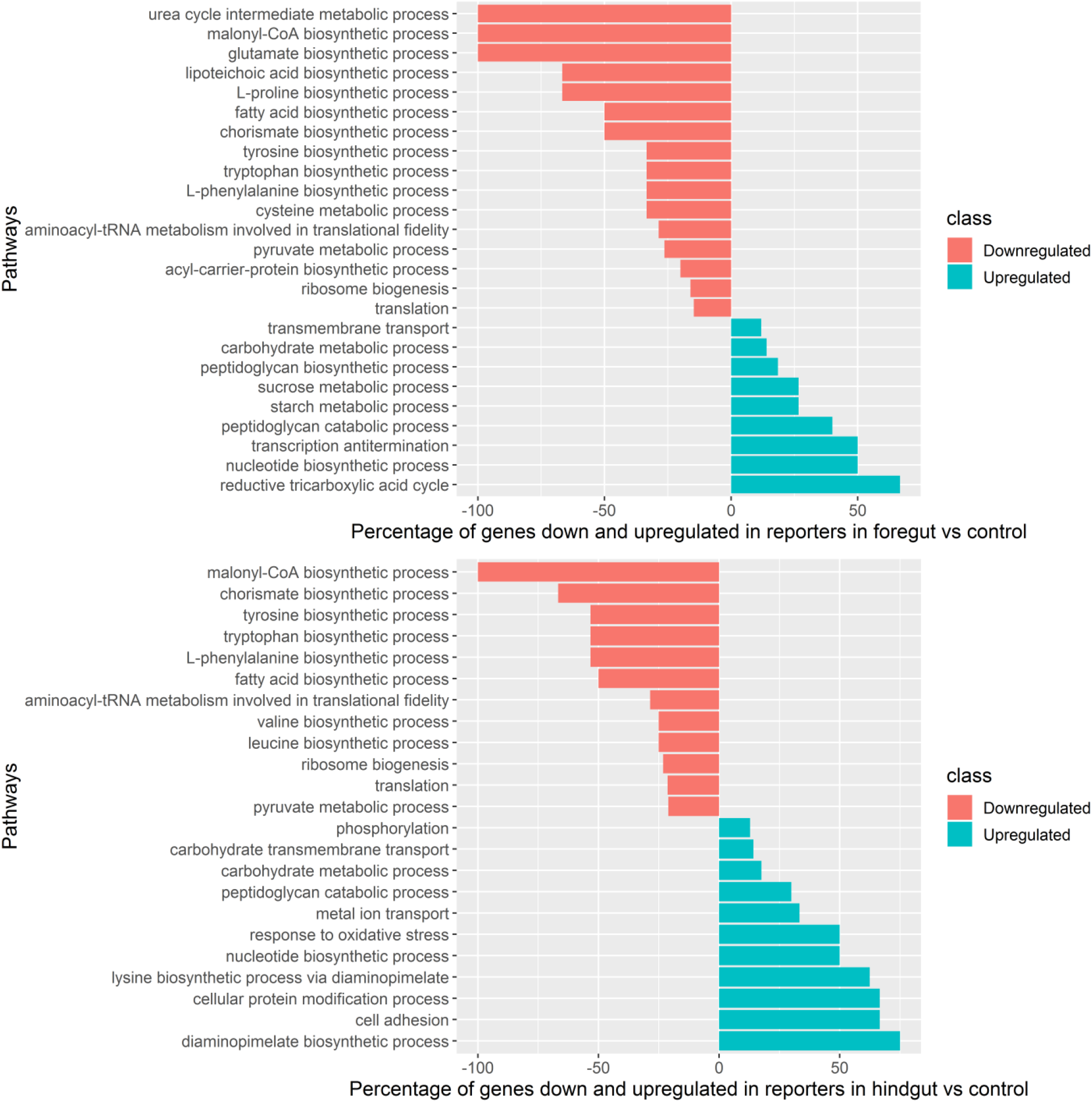
Summary of gene ontology classification in the category of biological processes. The graph shows both up- and downregulation of the assembled genes of *E. mundtii* obtained from foregut (a) and hindgut (b), as compared to genes of the control.

**Peptidoglycan turnover** is a sign of active cell division. Peptidoglycan biosynthetic and catabolic processes show upregulation in both fore- and hindguts. The N-acetylmuramoyl-L-alanine amidase enzyme in the foregut helps in cell separation during division (24). It also aids in cell motility and establishing a symbiotic association with the host (4).

Next task for *E. mundtii* would be to adhere to the host gut epithelium. Accordingly, the category **Cell adhesion** is enriched, which could indicate that the bacteria are involved in attaching and adhering to the host epithelium to prevent the flushing out of the host gut.

Aerobic respiration in insects and the electron transport chain in the mitochondria lead to reduction of oxygen to water and formation of ATP, resulting in the production of reactive oxygen species (ROS) at the gut membrane (25). Symbionts need to be able to respond to **oxidative stress** in order to live in the gut.

Certain metabolic pathways are worth mentioning:

**Nucleotide metabolism** seems to be a key aspect for bacterial colonization and adaptation to the host gut (26). Several enzymes required for purine and pyrimidine metabolism are upregulated.

An upregulation in **starch and sucrose metabolism** indicates that the symbionts are able to metabolize these complex sugars most readily because these are components of the larvae’s plant-based carbohydrate diet. The larvae show enriched alpha amylase activity, along with starch and sucrose metabolic processes in the foregut (27).

Not only the **synthesis of amino acids** such as phenylalanine, glutamate, tyrosine and tryptophan, but also **fatty acid biosynthesis** (shown by the downregulation of acetyl CoA carboxylase activity, malonyl CoA biosynthetic activity) and metabolism in general seem to be downregulated in the symbiont. There is a concomitant upregulation of **fatty acid degradation** when *E. mundtii* lives in the gut (Fig. S4). Perhaps by obtaining these by-products from the host, symbionts avoid the energy costs associated with these processes.

However, in the *E. mundtii* that reside in the hindgut, **lysine biosynthesis** via the diaminopimelate pathway is upregulated. Most likely this essential amino acid is provided to the host and may get reabsorbed in gut of *S. littoralis* (9) (Fig. 4(a, b)).

### 4. Survival strategies of *E. mundtii* in the gut of *S. littoralis*

The differentially expressed genes that were identified pertained to the adaptive strategies of *E. mundtii* in the fore- and hindguts of the larvae. We classified their strategies in three broad categories: extracellular interactions, stress responses and metabolism.

#### a) Extracellular interaction

Bacterial adherence to the gut tissue is the first step towards a successful colonization of bacteria in the host gut to prevent bacteria from being flushed out of the system and to survive the epithelial turnover (28).

Various well-characterized surface-associated proteins with conserved motifs and domains contribute to the attachment of *E. mundtii* to the gut epithelial tissue of the host. C-terminal conserved LPxTG motif (EMQU_1297: 26 and 40 folds in fore- and hindgut, a slight upregulation of fms3) and WXL domains (EMQU_0541: 30 and 8 folds in fore- and hindgut and EMQU_0539: 383 folds in foregut) are two such well-characterized and conserved parts of proteins that help bacteria to attach successfully to the host gut. Adhesion, the first step towards biofilm formation, is mediated by competition for nutrients. The *lysM* domain helping in biofilm formation (29) by surface attachment is upregulated (EMQU_0157: up to 3 folds in fore- and hindguts). The sticky matrix helps *Enterococcus mundtii* to deal with stress efficiently (28). The *agr* two-component systems may bring about quorum sensing in bacteria; when they come close together, the labor required to adapt to a new environment is divided among individuals. Levels of *agrA* are upregulated about 3-folds in the hind gut and for *agrB* about 5 and 8 folds in fore- and hindguts, respectively (30).

Chitin-binding proteins form a class of surface-associated proteins that provide adhesive properties to lactic acid bacteria so that these can adhere to the N-acetyl glucosamine component of chitin present in insects’ gut epithelial cells, especially the cells lining the midgut (31). Two of these proteins show levels as high as EMQU_0940: 47 and 138 folds and EMQU_1285: 25 and 69 folds, in fore- and hindguts, respectively (32).

Lipoproteins are placed at defined subcellular spaces formed by the plasma membrane. Their position is convenient to capture incoming nutrients or elements such as iron. Also, they have been shown to play roles in adhesion to host cells as well (33) (34). EMQU_0428 is upregulated by 5 and 4 folds in fore and hind gut respectively. EMQU_2743 is 7 folds upregulated in hind gut. Both are zinc transporter lipoproteins (Fig. 5, supplementary S5).

**Figure 5:**
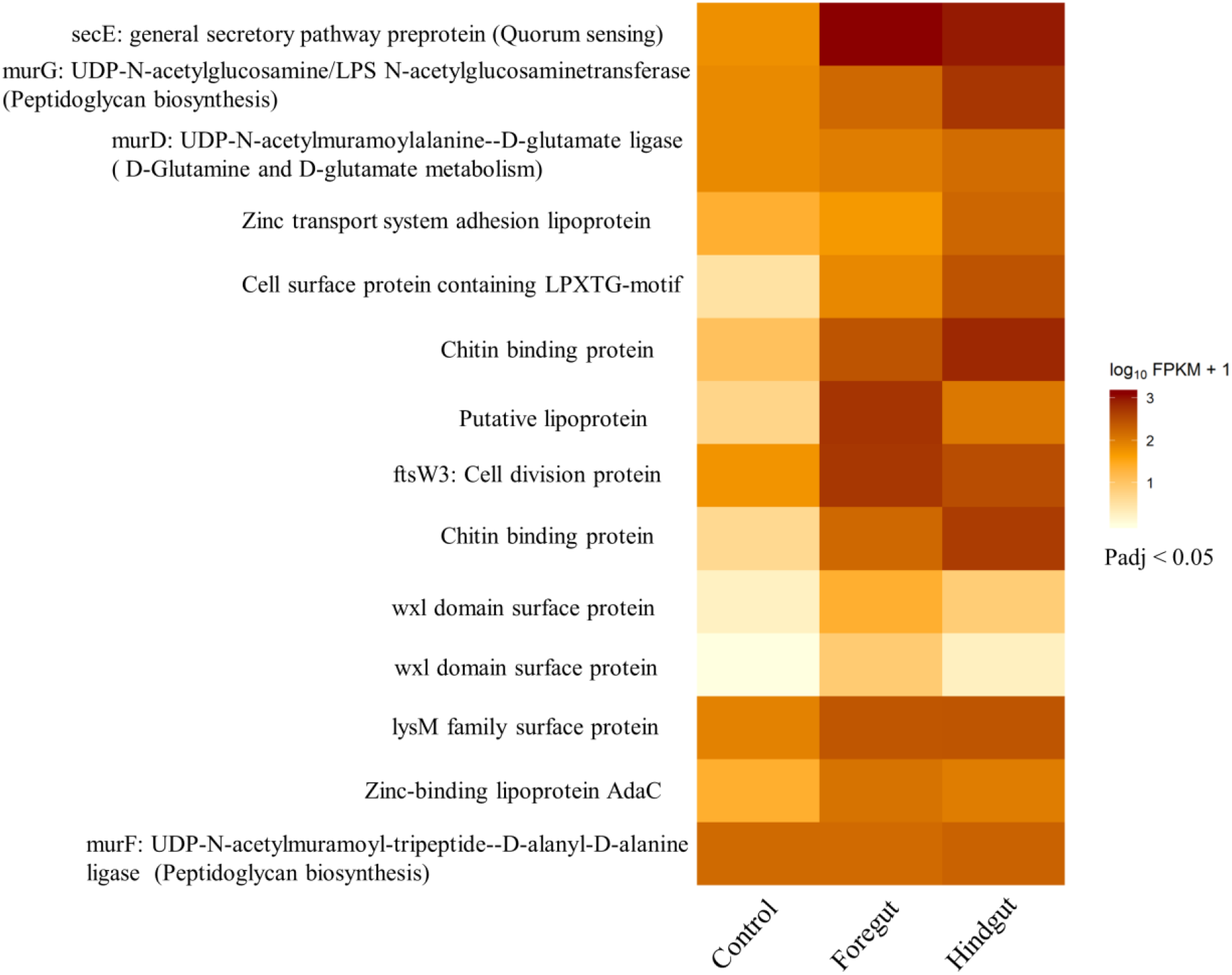
Heatmap showing the regulation of certain genes helping in the attachment of *E. mundtii,* when these bacteria are in the fore and hind guts of *Spodoptera littoralis*

#### b) Stress responses

Bacteria modulate their gene expression to abate the various stresses they encounter in the environment.

Quorum sensing in *E. mundtii* in the gut is mediated by the *agr* two-component system (35). *E. mundtii* residing in the insect gut are exposed to oxidative stress resulting from the host metabolism. Accordingly, they upregulate several antioxidant enzymes: superoxide dismutase (13 and 8 folds in fore- and hindguts, respectively), catalase (EMQU_0568: 4 and 10 folds in fore- and hindguts, respectively), NADH oxidase-peroxidase cycle (EMQU_0335, 0459, 1279; up to 4-folds in the hindgut), organic hydro peroxide resistance family protein (EMQU_1453: 6 folds in fore- and hindguts), and peptide-methionine (R)-S-oxide reductase (EMQU_0165: 3-folds in hindguts).

General stress proteins (*glsB*: 25 and 43 folds; *glsB1*: 11 and 8 folds; *gls33*: 7 and 19 folds, in fore- and hindguts, respectively) and universal stress proteins (USPs) (*uspA2*: 33 and 12 folds in fore- and hindguts, respectively) are upregulated in *E. mundtii* in response to environmental conditions such as salt, oxygen or oxidative stresses, toxic substances and nutrient starvation. The expression of USPs may depend on increased bacterial density, brought about by quorum sensing (36).

Intracellular trafficking, secretion, and vesicular transport include *secE* (19 and 16 folds in fore- and hindguts, respectively) needed for cell viability (37), and *virD4* components (31 and 30 folds in fore- and hindguts, respectively) of the type IV secretion system, which are all upregulated.

Repair proteins such as MutS (EMQU_2803) conferring DNA mismatch repair and its protection from oxidative stress are slightly upregulated). So is the *recA* (EMQU_2752: 2 folds in the foregut) operon that was shown to be upregulated during oxidative stress (38); *recF* for recombination repair (39), whose general role is maintenance of DNA is upregulated (2 and 3 folds in fore- and hindguts, respectively) (40).

DNA alkylation repair protein (AlkD) is upregulated 2 and 3 folds in fore- and hindguts, respectively (41). Also, RadA (3 and 7 folds in fore- and hindguts) and RadC (2 folds in fore- and hindguts) proteins helping in DNA repair and recombination show the same trend (42).

YafQ (EMQU_3002) and DNA damage-induced protein J (EMQU_3001, 25 and 4 folds in fore and hindguts, respectively) constitute a toxin-antitoxin system that plays a role in biofilm formation (43) (Fig. 6, supplementary S5).

**Figure 6:**
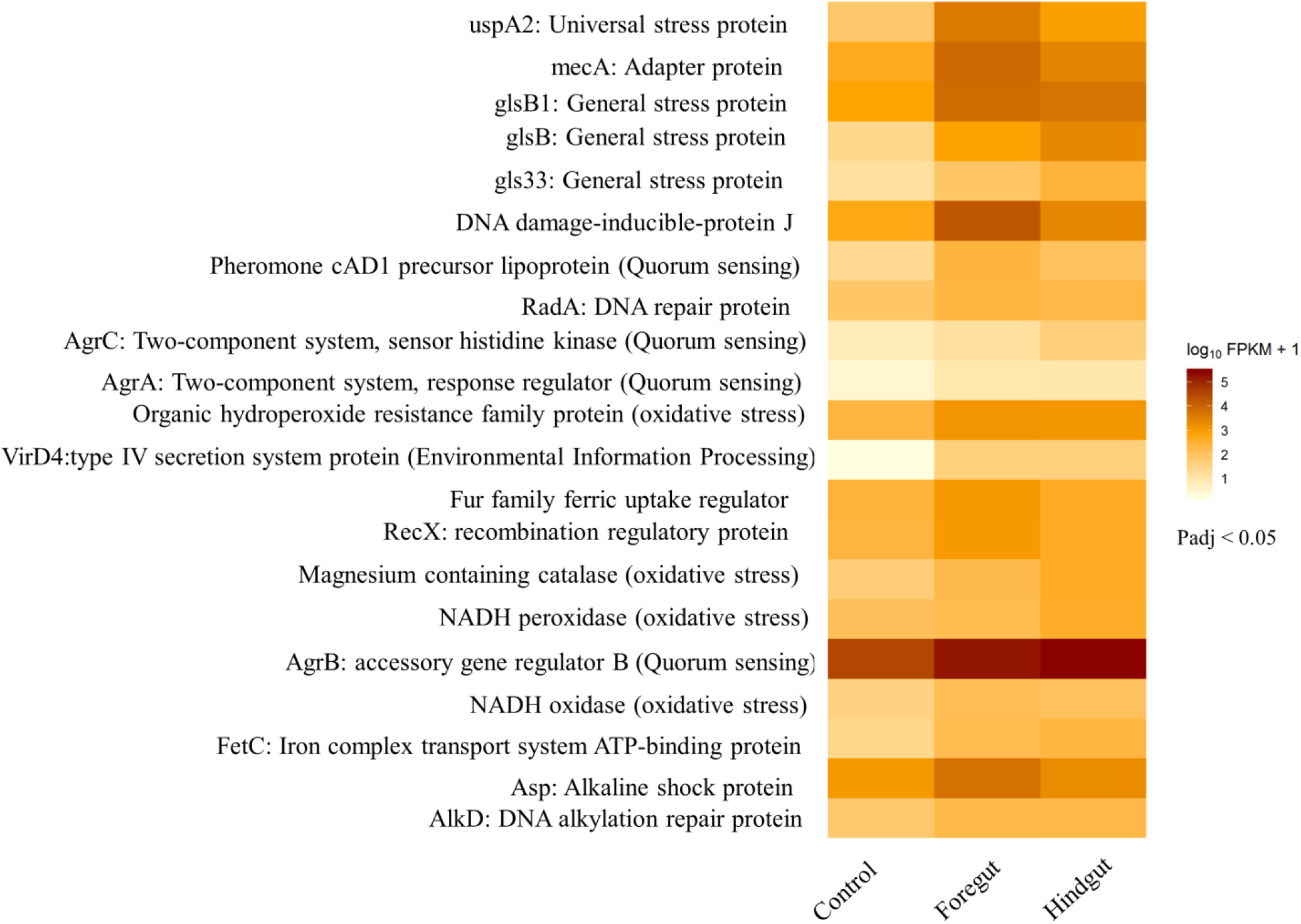
Heatmap showing the regulation of certain genes that help in the stress tolerance of *E. mundtii*, when these bacteria are in the fore and hind guts of *Spodoptera littoralis*

##### Iron homeostasis

Iron homeostasis in *E. mundtii* is important especially in environments that are iron depleted, owing to the compound 8-HQA. These bacteria have upregulated their *fetC* permease gene to increase their ferric uptake (7 folds in the foregut and 11 folds in the hindgut) and FUR family transcriptional regulator (EMQU_1067: 4 folds in the foregut) (FetC permease assists in siderophore-mediated iron uptake) (44). Adaptation that is mediated through FUR and iron uptake is common in iron-deprived environments (45) (Fig.6, supplementary S5).

##### Dealing with alkaline stress

The highly alkaline pH characteristic especially of the foregut of larvae is a challenge to bacteria in general but also to *E. mundtii* specifically. For example, alkaline pH has been proven to unwind the double helical structure of DNA (46). In addition, the alkaline stress protein, which has been reported to accumulate in the cell membrane of *Staphylococcus aureus* in alkaline conditions (47), shows high expression levels in *E. mundtii* living in the alkaline foregut (7 folds), while its expression decreases in the neutral conditions of the hindgut. In *Enterococcus faecalis,* several genes are differentially expressed in alkaline conditions; similar expression patterns characterize *Enterococcus mundtii,* if alkalinity is the only factor taken into consideration. For example, we found a downregulation of methionine transport and synthesis systems, Na^+^H^+^ antiporter (NhaC family, 1 fold downregulation), upregulation of adenosine and cytidine deaminases (upto 19 folds), purine and pyrimidine metabolism. The expression levels of Cation/H^+^ related antiporters (atp and ntp family proteins) are reduced under alkaline conditions (Fig. 7, supplementary S5, S6).

**Figure 7:**
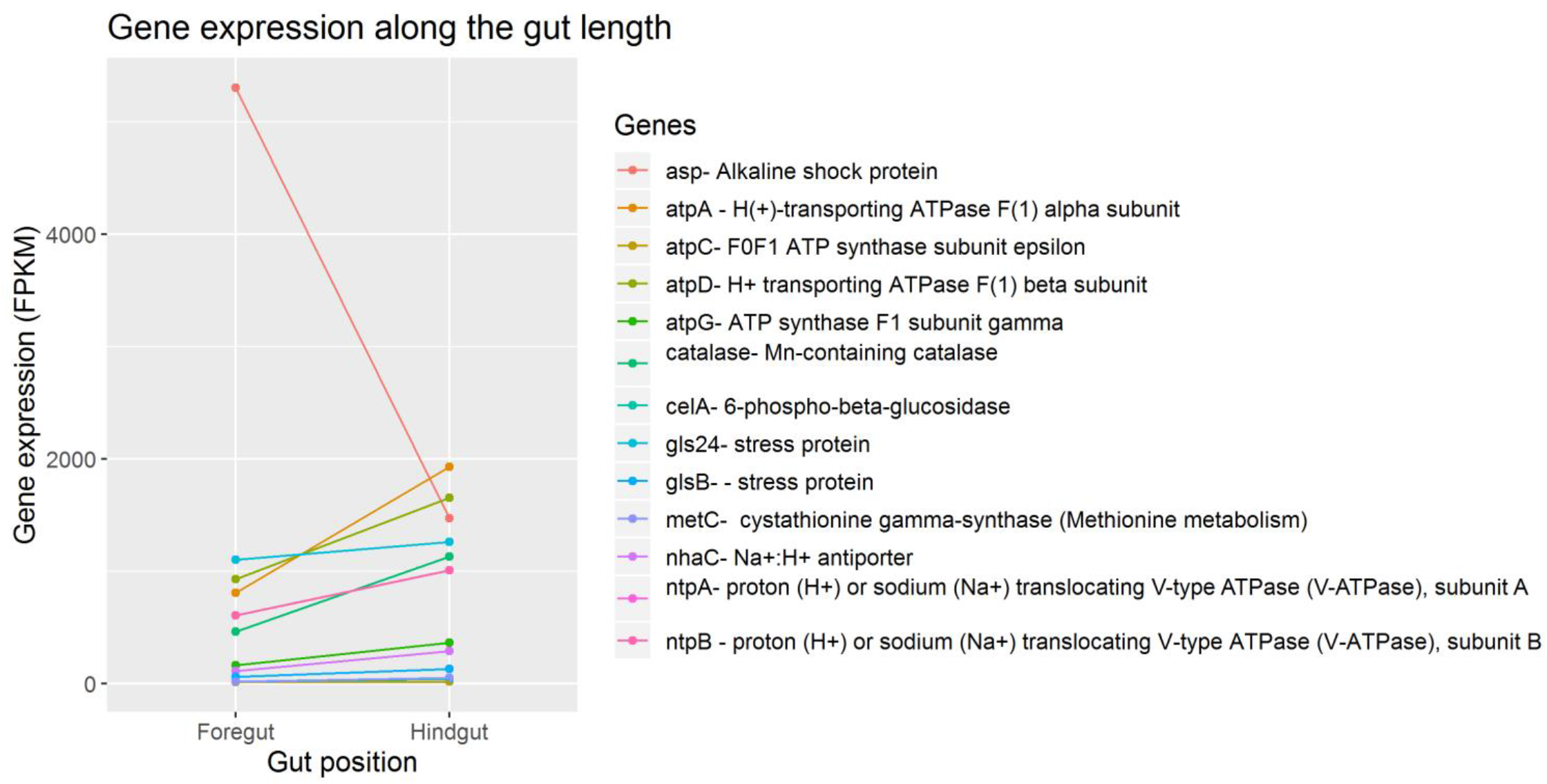
Graph showing the regulation of certain pH-related genes in *E. mundtii* living in the fore and hind guts of *S. littoralis* larval gut.

#### c) Metabolism

Facultative anaerobes can switch between respiration and fermentation, based on oxygen availability. Most glycolytic genes–for example, glucokinase (*glcK*), 1-phosphofructo kinase (*fruK*), 6-phospho-beta-glucosidase (*bglP, bglB, bglG*) phosphofructokinase A (*pfkA*) and glucose-6-phosphate isomerase-in the *E. mundtii* dwelling in the gut do not show much change in expression compared to those growing under control conditions, suggesting that glycolysis has not been stopped completely. The same trend holds true for pyruvate dehydrogenase for entering the citric acid cycle in aerobic condition, along with lactate dehydrogenase (*ldhA* EMQU_2453). The protein that stimulates the sugar fermentation (SfsA-EMQU_0871) under anaerobic conditions is upregulated 10 and 7 folds in fore- and hindguts, respectively. Some alcohol dehydrogenases are upregulated to convert acetaldehyde to ethanol in the fermentative pathway (EMQU_1129, 1 fold in hindgut; EMQU_ 0525, 5 folds in fore- and hindguts, and EMQU_0315, 3 and 4 folds in fore- and hindguts). The acetyl CoA produced by pyruvate dehydrogenase does not significantly contribute to the production of fatty acids and amino acids, because both pathways are downregulated (Fig. S4).

Phosphotransferase systems (PTSs) for taking up alternative source of sugars such as sucrose, ascorbate, mannose and, most importantly, cellobiose, are upregulated in *E. mundtii* in both fore- and hindguts. Cellobiose comes from the plant products on which the host is fed. There are at least 8 PTS cellobiose transporters upregulated with fold-changes as high as 39 and 42 in the fore- and hindguts, respectively. Ascorbate is mostly taken up in the hindgut. On the other hand, fructose and lactose do not seem to be a popular source of energy.

There is a dramatic increase in starch and sucrose metabolism brought about by an increase in the sucrose-specific PTS transporter (EMQU_2136: 1 and 5 folds in fore- and hindguts, respectively) and sucrose 6-phosphate dehydrogenase (*scrB*: 2 folds in the hindgut); and the alpha-amylase enzyme neopullanase (EMQU_1435: 52 and 30 folds in the fore- and hindguts, respectively).

There is an increased metabolism and transport of nucleotides in *E. mundtii* living in the gut (26).

Regarding glycerol metabolism, the *glpF* gene required for glycerol uptake is downregulated (4 folds in the foregut), whereas the ones for metabolism-*glpO, dhaKL, glpQ* are still expressed, suggesting an alternate way for these bacteria to obtain glycerol (48) (Fig.7, supplementary S5).

**Figure 8:**
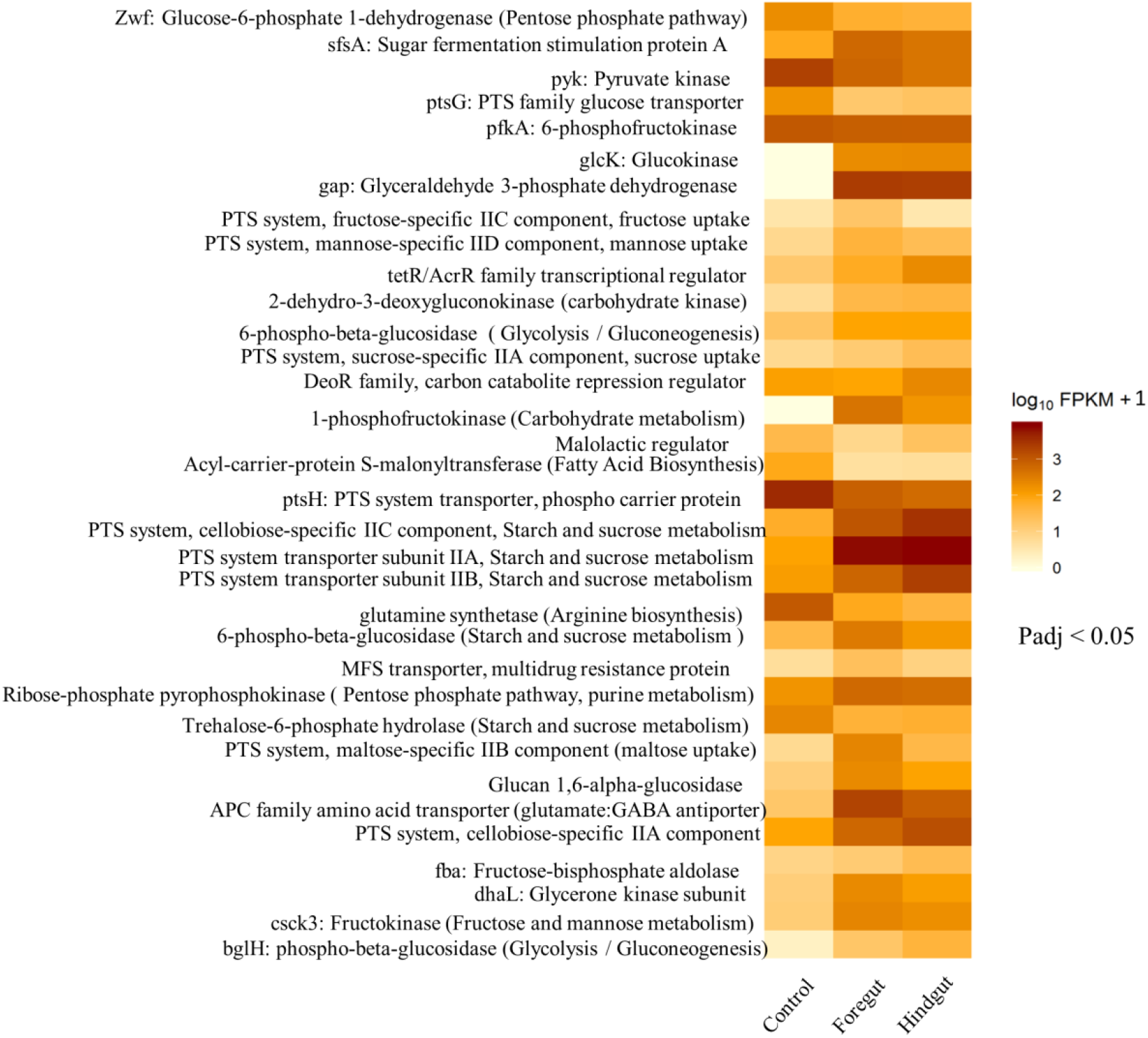
Heatmap showing the regulation of some genes in *E. mundtii* are involved in metabolism when they are in the fore and hind guts of *Spodoptera littoralis*

## Discussion

This work focuses on the survival strategies of *Enterococcus mundtii* in the gut of *Spodoptera littoralis*, an environment threatened by stressful conditions, namely high pH, low iron content and oxidative stress. This makes it a good system to study adaptation by the symbionts in the larval gut. This motivated us to send down the larval guts, GFP-tagged reporter *E. mundtii*, which is a dominant bacterium (18), in order to study how its mechanisms of adaptations to the new environment. The fed fluorescent bacteria were later retrieved from the fore and hindguts of the larvae using flow cytometry. Care was taken to halt any kind of metabolic changes that might have occurred between the individual experimental steps of larval dissection and FACS sorting, using RNAlater and RNAprotect reagents. Comparing the gene expression profiles of these retrieved reporters with those of *E. mundtii* grown under optimal culture conditions, we were able to obtain a snapshot of the genes and the pathways that help these symbionts to survive and adapt to the gut of *S. littoralis* larvae. The transcriptional changes found in these bacteria are an amalgamated result of all these factors.

We managed to obtain a real-time scenario of how *E. mundtii* is abating stress and colonizing its host gut (Fig. 9).

**Figure 9:**
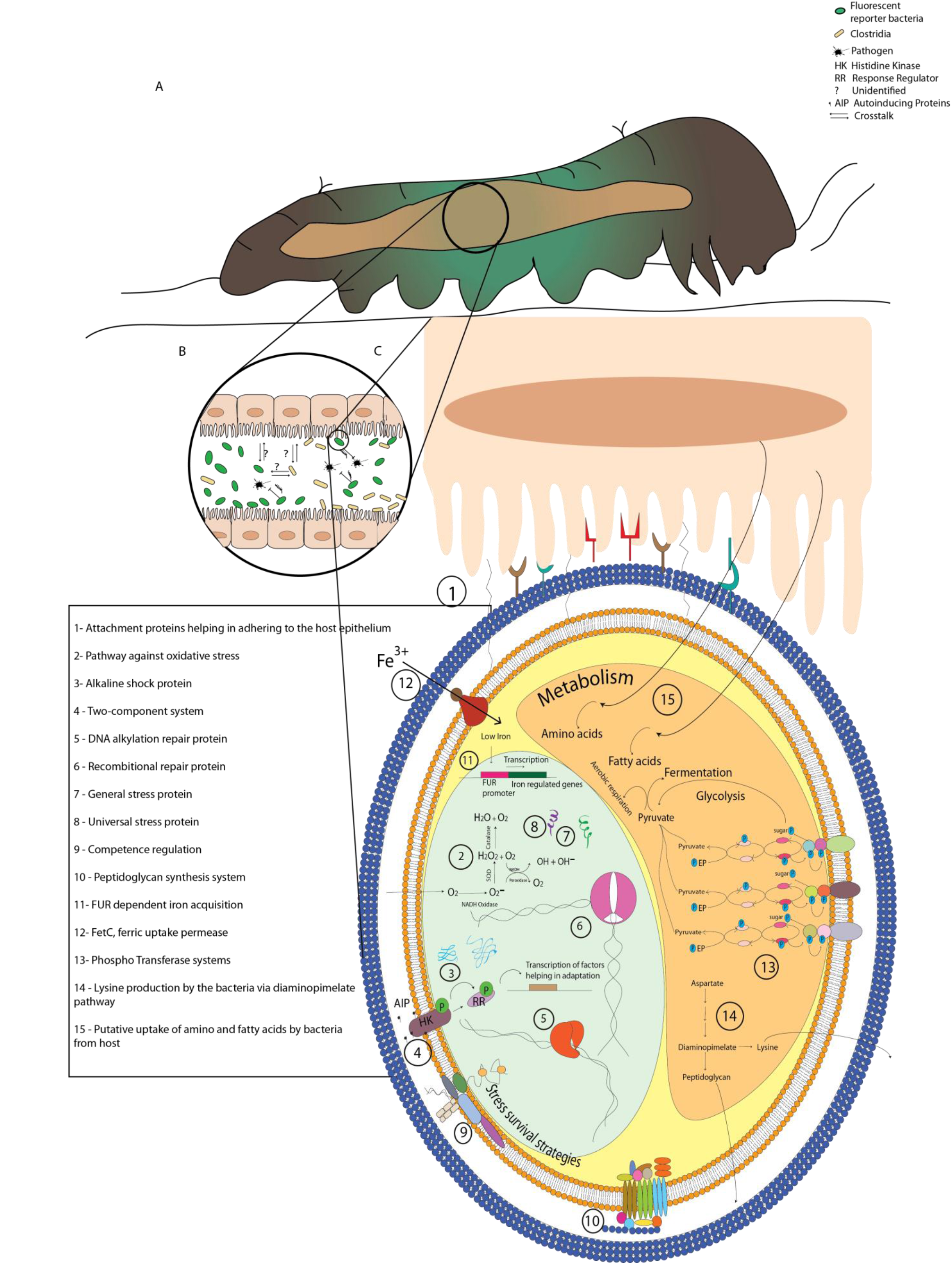
A snapshot of interactions between *S. littoralis* and its resident gut symbiont, *E. mundtii*. (A) An illustration of *Spodoptera littoralis* with its longitudinal gut (B) *E. mundtii* dominates in the gut along with Clostridia and keeps pathogens at bay by producing mundticiin KS. Unknown interactions occur among these two symbionts and the host gut. (C) Pathways and stress survival strategies of *E. mundtii* in the gut of *S. littoralis*.

The ability to adhere to the host gut epithelium is an important prerequisite for the bacteria to successfully colonize and establish a host-microbe association. Most of the adhesive properties of lactobacilli are accounted for by their cell surface properties (49).

Our transcriptomic analyses show that *E. mundtii* attaches well to the host surface, as was also seen earlier by FISH imaging before (5). This is brought about by a set of adhering proteins. LPxTG is a sortase-dependent site for anchoring proteins, covalently attached to the peptidoglycan (50), (51). Lipid-anchored proteins or lipoproteins constitute another class of covalently associated adhesion proteins (49), which shows upregulation in *E. mundtii*. Wxl domains and LysM, or lysine-dependent motif binding to the peptidoglycan, form non-covalent associations with the peptidoglycan (52), (53), (52). The occurrence of such associations was proven in *Enterococcus fecalis* (54). Chitin is a major part of the peritrophic matrix, which in turn lines the midgut epithelium of the host (55), (56). Chitin-binding proteins in *E. mundtii* also promote adherence to the host gut. Several bacteria such as *L. monocytogenes*, adherent *E. coli* and *V. cholerae* were found to initiate their adhesive process using their chitin-binding proteins in the host gut (57).

*E. mundtii* dwelling in the gut employ various strategies to survive adverse conditions, upregulating general and universal stress proteins to survive heat, pH and oxidative stress. Mechanisms to abate oxidative stress are a necessity for organisms living in an actively metabolizing environment. Reactive oxygen species result from reduction of oxygen. Thereafter, the dismutation product of the superoxide anion (O_2_^-^) is hydrogen peroxide (H_2_O_2_) (58). O_2_^-^ and H_2_O_2_, along with the hydroxyl radical, are potent oxidants that can abstract electrons from DNA, proteins, lipids, other macromolecules, causing damage to the invading or residing symbionts (59). Lactobacilli employ enzymes such as NADH oxidase/peroxidase, superoxide dismutase and manganese-dependent catalase to counteract ROS (60).

Several bacteria have been found to have general and universal stress proteins, aiding their adaptation to stresses such as temperature, oxidative, nutrient starvation and toxic agents (61).

The first reported universal stress protein in *E. coli* was found in fungi, archaea, plants and even flies (62). In *Burkholderi aglumae*, universal stress proteins genes regulated by quorum sensing (61). Confronted with stress, *E. mundtii* seems capable of behaving like a multicellular organism. The bacteria rely on quorum sensing as a survival strategy, aggregating on the host epithelia and forming a biofilm in the host gut. That *agrABCD* forms a two-component system and brings about quorum sensing has already been established in fellow fermicute *Staphylococcus aureus* and *Streptococcus pneumoniae* (63). The bacterial adherence properties of *E. mundtii* may help form a biofilm layer on the gut wall. Thus, these two inter-related phenomena, quorum sensing and biofilm formation, and help bacteria to adapt to altered environments.

As discussed, 8-HQA produced by the larvae is an iron chelator, and the symbionts must employ mechanisms to survive in an iron-depleted environment. The FetC iron complex transport permease and FUR family of transcriptional regulators seem to have similar goals. FetC was found to be involved in iron homeostasis in *Apergillus fumigates* (64). FUR-dependent iron-acquisition system was upregulated when *Clostridium difficile* tried to infect hamsters in iron-depleted conditions (65).

Owing to the alkaline environment of the foregut, the *E. mundtii* living there highly express alkaline shock proteins as protection against alkaline stress. Such was also the case in *Staphylococcus aureus* (66). The alkaline shock protein helps the bacteria to adapt to extreme stress conditions (67). All the F and V-type ATPases (atp, ntp genes), (Fig. 5b) show downregulation in an alkaline environment. At high alkaline pH, the proton motif force decreases drastically, and lowers the activity of H+ ATPases. These ATPases are related to energy production and conversion. A similar trend was found in *E. fecalis* growing in an alkaline environment (68).

As facultative anaerobes, *Enterococcus mundtii* has the tendency to switch to fermentation inside the host gut, although aerobic respiration is not completely stopped, since the genes for both are upregulated (fig 9). Oxygen may only be found in the near vicinity of the host gut surface, 50 μm distance onwards, for the facultative anaerobes, since there is none in the inner layers of the anaerobic gut wall (5). This tells us about the low oxygen levels in the gut lumen of most insects (69). Pathway analysis clearly shows that conditions are favorable for starch and sucrose uptake through PTS transporters and metabolism. This is accounted for by the white-bean-based artificial diet that the host is fed on. The PTS transporters are immensely helpful for bacteria in general to survive environments with different levels of sugar (70). Nucleotide metabolism shows enrichment owing to the fact that the bacteria are striving to colonize the gut of *S. littoralis*. Previous studies with mice models showed that *E. coli* enriched their purine and pyrimidine metabolism in order to colonize the mice intestines (26). Although *E. mundtii* likes to save energy expenditure for its fatty acid and amino acid metabolism, their lysine metabolism is upregulated by bacteria living in the hindgut (9). Whether *S. littoralis* is obtaining lysine from their symbiotic *E. mundtii*, is a matter of further research. Pathway analysis shows enrichment for lysine synthesis via the diaminopimelate pathway. Diaminopimelate also plays roles in peptidoglycan synthesis (71).

High-throughput transcriptome sequencing from tiny quantities of starting material has provided insights into the strategies used by *E. mundtii* to survive the gut of *S. littoralis,* as real-time information coming straight from the gut. The method optimized in this work can be put to effective use for studying interactions between a pair of any host and its symbiont. For example, the future prospects of this work include how 8-HQA is acting as a deciding factor in manipulating the bacterial community. A similar methodology will allow us to introduce fluorescently tagged bacteria into the insect guts that have been knocked out of the 8-HQA producing gene, or any other gene that seems to be interesting for a symbiont’s survival. Quite similar to our method, the behavior of the retrieved bacteria studied with omics can tell us how they act in such varying conditions. This way, we can delve deeper into interaction studies to allow us a better understanding of mechanisms of survival and exchange occurring among communicating partners in the environment.

## Materials and methods

### Maintenance of eggs and larvae

The eggs of *Spodoptera littoralis* were obtained from Syngenta Crop Protection Munchwielen AG (Munchwielen, Switzerland). Eggs were hatched at 14^°^ C and the larvae were maintained at 24^°^ C in an alternate 16 hours light period and 8 hours dark period. Larvae were reared on an agar-based artificial diet containing white beans, as described by Maffei et al (72).

### Bacterial strain

A fluorescent strain of *E. mundtii* KD251 (isolated from the gut of *S. littoralis* in the Department of Bioorganic Chemistry) was constructed by transforming a GFP containing expression vector pTRKH3-ermGFP as described (18). This strain was grown in Todd-Hewitt Bouillon, THB (Roth, Karlsruhe, Germany) medium for both broth and 1.5% agar (Roth, Karlsruhe, Germany), and in the presence of 5 μg ml^-1^ of erythromycin (Acros Organics, New Jersey, USA). The strain was preserved as a glycerol stock at −80^°^ C.

### Introduction of the reporter bacteria into the insect microbiome

A stationary phase culture of fluorescent reporter *E. mundtii* in THB broth containing 5 μg ml^-1^ of erythromycin was grown till mid-log phase with OD_600_ ~ 0.5-0.6 at 37°C with shaking at 220 rpm. The culture was pelleted at 5000 x g for 10 minutes at 4°C and resuspended in distilled water. First-instar *S. littoralis* larvae (n = 120) were fed small cubes of artificial diet supplemented with two antibiotics, ampicillin (5.75 μgml^-1^) (EMD Millipore corp., Billerica,

MA, USA) and erythromycin (9.6 μgml^-1^) for 3 days, to reduce the already existing bacterial load, before being fed with (at second instar) 100 μl from the 1:10 dilution broth (~10^^10^ cells) containing fluorescent *E. mundtii* as described (18). These larvae were allowed to grow until fifth instar, until sample preparation for FACS.

### Sample preparation for FACS

A total of 30 fifth-instar larvae for each gut region, foregut and hindgut were dissected with sterile forceps and scissors in a sterile clean bench. Following dissection, the gut tissues were immediately submerged in 10 ml of RNAlater solution (Invitrogen, Vilnius, Lithuania). Tissues submerged in RNAlater solution were mixed with 2 ml of 6% (w/v) betaine (Sigma Aldrich, St. Louis, MO, USA) and placed on ice prior to crushing with mortar and pestle until gut homogenates were formed. Thereafter, fluorescent *E. mundtii* were separated from the intestinal debris by filtration through 40 μm pore-size cell strainers (Falcon, NY, USA). The filtrates were then separated into aliquots of 600 μl each and kept at −80°C for the sorting experiment.

### Cell sorting by FACS

The gut homogenates were analyzed using BD FACSAria^™^ Fusion Cell Sorter (Becton Dickinson, Heidelberg, Germany). It utilizes an ion laser emitting a 488 nm wavelength, and a 502 long pass filter, followed by a 530/30 band pass filter. The green fluorescent protein emits light with a peak wavelength of 530 nm. The cells were sorted at a flow rate ranging between 10 μlmin^-1^–80 μlmin^-1^. The sorting was done in a single-cell mode and the sorted cells were collected in 5 ml sterile Polypropylene round-bottom tubes (Falcon, Mexico). The cells were collected for a period of 3 hours which corresponded to an acquisition of 6000-7000 events/sec. The flow cytometry grade of PBS buffer (Thermo Fischer, Wilmington, USA) at pH of 7.4 was used as the sheath fluid. A total of ~ 250, 000 cells were sorted from each sample, into 1 ml of RNA Protect solution (Qiagen, Hilden, Germany)

### RNA extraction and sequencing

As controls, *E. mundtii* broth cultures (10 ml, *n* = 3) were grown to exponential growth (OD_600_ ~ 0.5-0.6) and centrifuged at 5000 x g for 15 min at 4°C to pellet the bacterial cells. Bacterial cells were washed once with sterile phosphate-buffered saline (PBS) and resuspended with the same buffer at a concentration of approximately 10^10^ CFU ml^-1^. The FACS-sorted fluorescent bacterial cells (~ 250,000) each from foregut and hindgut were pelleted by centrifugation at 5000 x g for 10 min at 4°C, leaving insect cell debris in the supernatant. The foregut, hindgut and *E. mundtii* cultures were each represented by three biological replicates (*n* = 3). RNAlater was removed from the sorted cells prior to RNA isolation and total RNA was isolated from the pelleted cells using the RNeasy mini kit (Qiagen, Hilden, Germany) following the manufacturer’s instructions with some modifications. Pelleted bacterial cells were lysed enzymatically for 15 min at 37°C (enzymatic mix: 1X TE buffer, pH 8 (Applichem GmbH, Darmstadt, Germany), pH 8.0, 5 μg ml^-1^ lysozyme (Sigma Aldrich, St. Louis, MO,USA) and 50 Uml^-1^mutanolysin (Sigma Aldrich, St. Louis, MO,USA)). All samples were DNase-treated with on-column DNase digestion per the manufacturer’s protocol prior to RNA isolation. The concentration of total RNA of controls was diluted to match the bacterial concentration at single cell level. RNA was further cleaned and concentrated using Concentrator kit (Zymo Research, USA) yielding about 12 μl in final volume (~10 ng). The purified RNA was linearly amplified using MessageAmp II bacterial RNA amplification kit (Invitrogen, Vilnius, Lithuania) using 10 ng of total RNA following the manufacturer’s instructions. The amplified RNA (aRNA) was concentrated by precipitation with 5M ammonium acetate.

The quality and quantity of the total RNA was measured with a NanoDrop One Spectrophotometer (Thermo Scientific, Wilmington, USA) and an Agilent 2100 Bioanalyzer (Agilent Technologies, Palo Alto, CA, USA). RNA samples were sent to the Max Planck Genome Centre in Cologne for RNA sequencing. A total of 0.3 μg −1 μg of amplified RNA was used for cDNA library preparation using the Ultra-Low Input RNA kit following the Illumina protocol at the Max Planck Genome Centre, Cologne. Sequencing was carried out on the HiSeq 2500 sequencer at Cologne and a total of approximately 10 million paired-end reads (2 x 150 bp) were generated for each sample.

### RNA-seq data analysis

FastQC was done for an initial quality analysis of the reads. Analysis of the reads, including trimming of adapters and differential gene expression analysis was done on LINUX-based Command line interface, following the Tuxedo protocol (73). The adapters were trimmed using Trimmomatic 0.36, trimmed reads were assembled using Tophat 2.1.0, and mapped to the genome of *Enterococcus mundtii* QU25 (23) using Cufflinks 2.2.0. The read counts were normalized with FPKM (Fragments of Kilobase of transcripts per Million mapped reads) (supplementary S6) and assemblies were merged using Cuffmerge. Cuffdiff was used to compute the differentially expressed genes between *E. mundtii* from the larval gut and *E. mundtii* grown *in vitro.* Based on homology to protein families, the proteins that were predicted for *E. mundtii* were categorized under Gene Ontology terms (http://geneontology.org). The genes were also mapped to the KEGG database to predict the pathways (supplementary). The results of differentially expressed genes were visualized using R-package CummeRbund 2.0, on R version 3.3.3 (2017-03-06). This R-package generated all the plots: dendrograms, volcano plots and histograms. The PCA plot was made using Statistics Toolbox of Matlab 2007a. A fold-change of ≥ 2 was has been used as a threshold to analyze the differentially expressed genes.

## Supporting information

Genes and fold changes

Genes and FPKM values

Overlap of differentially regulated E. mundtii genes in fore and hind guts of the host

Supplementary sheet 1

## Acknowledgements

This work was funded by the Max Planck Society and Jena School for Microbial Communications. Also, we thank Angelika Berg and Andrea Lehr for technical assistance and Emily Wheeler for editorial assistance.

