## Supplementary sheet 1 for "Survival strategies of *Enterococcus mundtii* in the gut of *Spodoptera littoralis*: a live report"

Supplementary material

**S1**: Sorting profiles of *Enterococcus mundtii*-pTRKH3 from (a) control (without GFP), (b) foregut homogenate and (c) hindgut homogenate. 250,000 fluorescent cells were retrieved from each biological replicate of foregut, hindgut and culture-grown bacteria.

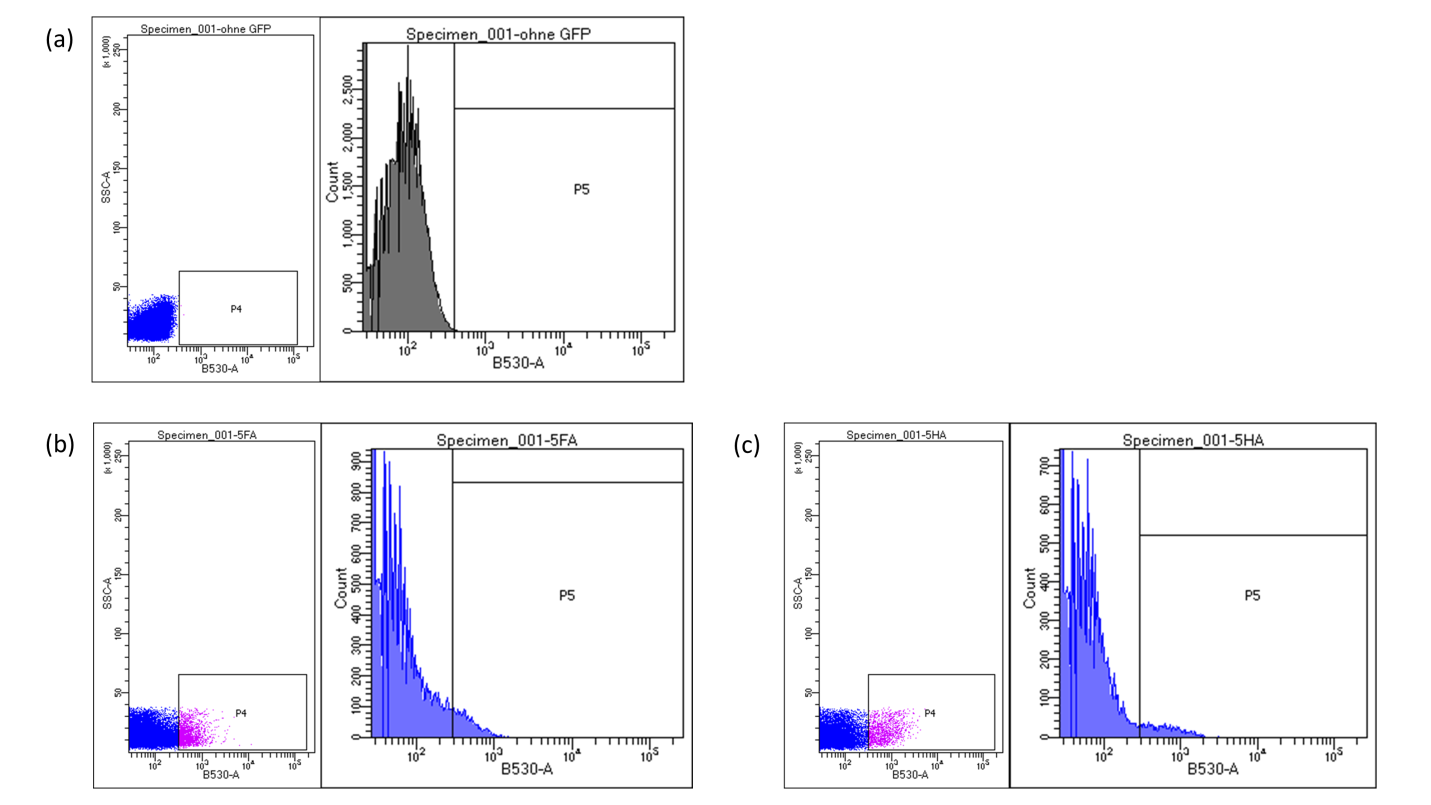

**Figure S1:** Sorting profiles of *Enterococcus mundtii*-pTRKH3 from (a) control (without GFP), (b) foregut homogenate and (c) hindgut homogenate. P4 and P5 correspond to the density of fluorescent cells detected by the scatters, and the cells that are actually sorted, respectively. The purple dots correspond to the GFP-containing *Enterococcus mundtii* that was sorted using the flow cytometer.

**S2**: Table showing the mapping percentages of the reads obtained from Enterococcus mundtii living in the foregut, hind gut of *Spodoptera littoralis* larvae, and the same bacteria grown *in vitro.* The mapping was done against the whole genome of *Enterococcus mundtii* QU25

| Sample | Replicates | # of reads from Trimmomatic; after adapter trimming | % alignment by Tophat |
| --- | --- | --- | --- |
| Foregut | F1 | 13949493 | 17.55 |
|  | F2 | 10562548 | 58.80 |
|  | F3 | 10623050 | 73.40 |
| Hindgut | H1 | 9491522 | 27.25 |
|  | H2 | 9161970 | 47.10 |
|  | H3 | 10187317 | 55.50 |
| Control | C1 | 9386723 | 41.55 |
|  | C2 | 9854743 | 38.70 |
|  | C3 | 9733545 | 48.25 |

**S3:** Distribution of the transcriptomic profiles of *E. mundtii* obtained from the foregut (FG), hindgut (HG) of *S. littoralis* and the ones growing *in vitro* control conditions (FL)

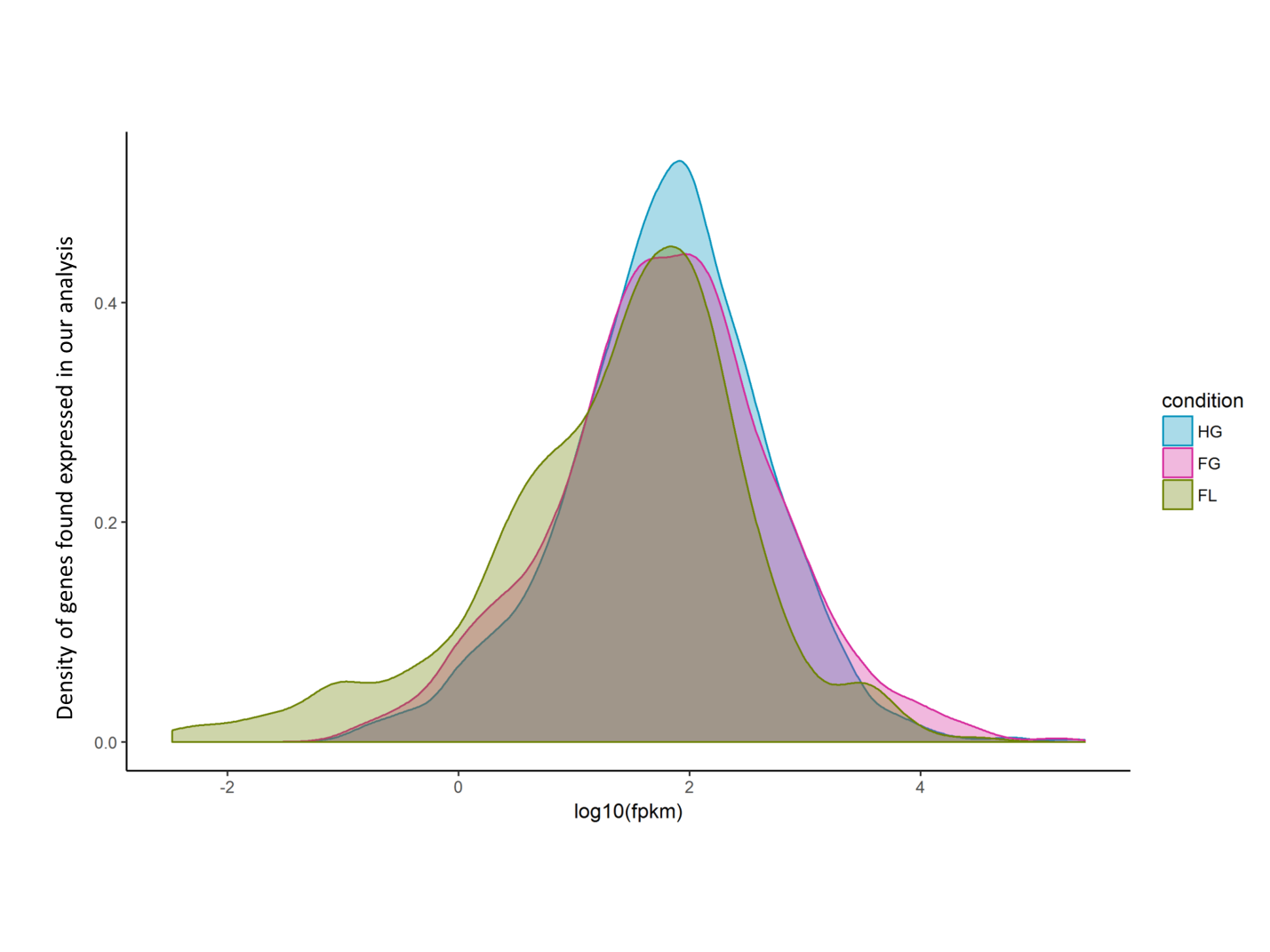

**Figure S3**: Density plot showing the distribution of the differentially expressed genes in foregut (FG), hindgut (HG) or control (FL)

**S4**: Annotations based on KEGG database

Annotations using the Kegg database show a similar trend. 938 differentially expressed genes in *E. mundtii* obtained from the gut were mapped to Kegg Pathways to give them a biological significance. 44 and 29 pathways were upregulated in foregut and hindgut respectively, whereas, 52 and 46 were downregulated. The ones that are significantly enriched are shown in figure (S4).

Degradation of aromatic compounds by the *E. mundtii* could be occurring due to the lignin content in the plant diet. The enrichment of starch and sugar metabolism remains similar.

Enrichment in napthalene degradation and chloroalkane and chloroalkene degradation mirror the high activity of alcohol dehydrogenases, which carries out steps in bacterial glycolysis and fermentation.

There are marked downregulations in several amino acid and fatty acid biosynthetic pathways, biosynthesis of antibiotics, propionate metabolism and biosynthesis of secondary metabolites; although the hindgut dwelling *E. mundtii* seem to biosynthesize lysine. It is quite possible that the lysine produced in the hindgut gets reabsorbed by the larval system through their hindgut intestinal cells.

S4 (a, b).
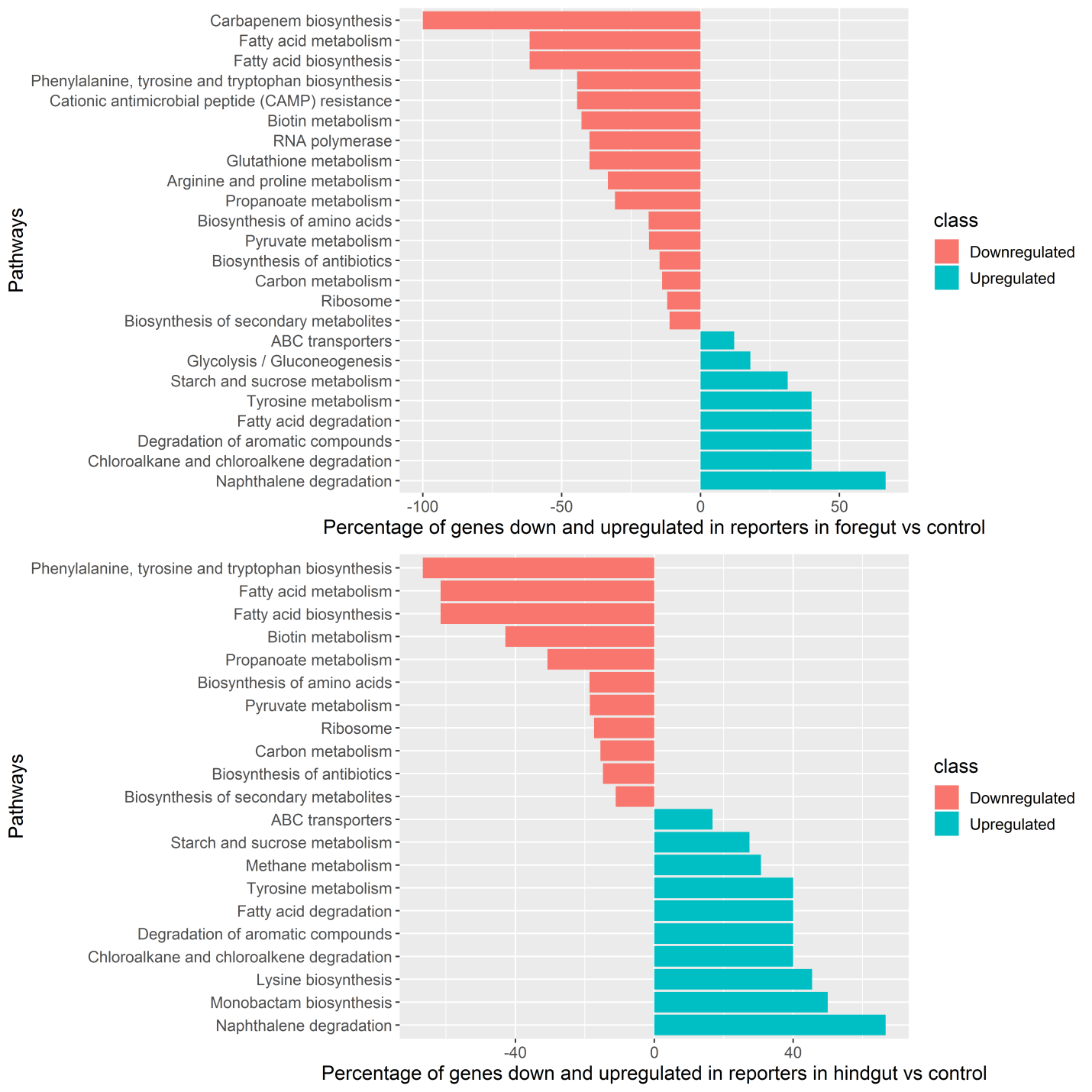

**Figure S4**: Kegg Orthology classification of assembled unigenes, annotated from transcriptomic information given by *E. mundtii* obtained from fore (a) and hindguts (b) respectively. The graphs show both up and down- regulation of the assembled genes, as compared to control.

**S5** (Excel sheet):

All the differentially expressed genes in Enterococcus mundtii when they are dwelling in:

Sheet 1: Fore gut vs Control

Sheet 2: Hind gut vs Control

Sheet 3: Hindgut vs Foregut

Sheet 4: Fold changes of selected genes of *E. mundtii*, pertaining to extracellular attachment, stress survival mechanisms and metabolism, when they are living in the fore and hind guts of *S. littoralis*.

**S6** (Excel sheet):

FPKM values of all the differentially regulated genes in *E. mundtii*, when they are living in the fore and hind guts of *S. littoralis.* All three replicates of foregut (FG), hindgut (HG) and control samples are shown.

**S7** (Excel sheet):

Genes that are up regulated in *Enterococcus mundtii* when they are dwelling in both fore and hindguts of *Spodoptera littoralis.*
